# ORBIT: a new paradigm for genetic engineering of mycobacterial chromosomes

**DOI:** 10.1101/249292

**Authors:** Kenan C. Murphy, Samantha J. Nelson, Subhalaxmi Nambi, Kadamba Papavinasasundaram, Christina E. Baer, Christopher M. Sassetti

**Affiliations:** Department of Microbiology and Physiological Systems, University of Massachusetts Medical School, Worcester, MA 01655

## Abstract

Current methods for genome engineering in mycobacteria rely on relatively inefficient recombination systems that require the laborious construction of a long double-stranded DNA substrate for each desired modification. We combined two efficient recombination systems to produce a versatile method for high-throughput chromosomal engineering that obviates the need for the preparation of double-stranded DNA recombination substrates. A synthetic “targeting oligonucleotide” is incorporated into the chromosome via homologous recombination mediated by the phage Che9c RecT annelase. This oligo contains a site-specific recombination site for the directional Bxb1 integrase (Int), which allows the simultaneous integration of a “payload plasmid” that contains a cognate recombination site and selectable marker. The targeting oligo and payload plasmid are co-transformed into a RecT‐ and Int-expressing strain, and drug-resistant homologous recombinants are selected in a single step. A library of reusable target-independent payload plasmids is available to generate knockouts and promoter replacements, or to fuse the C-terminal-encoding regions of target genes with tags of various functionalities. This new system is called ORBIT (Oligo-mediated Recombineering followed by Bxb1 Integrase Targeting) and is ideally suited for the creation of libraries consisting of large numbers of deletions, insertions or fusions in a target bacterium. We demonstrate the utility of ORBIT by the construction of insertions or deletions in over 100 genes in *M. tuberculosis* and *M. smegmatis*. The report describes the first genetic engineering technique for making selectable chromosomal fusions and deletions that does not require the construction of target‐ or modification-specific double-stranded DNA recombination substrates.

## INTRODUCTION

Following the use of laborious plasmid co-integration schemes that dominated the early days of gene replacement in bacteria, recent advances in genetic engineering of bacterial strains have largely relied on phage recombination systems, both site-specific and homologous. For example, it was recognized in 1994 that the use of non-replicating plasmids containing phage lambda *attP* sites could be incorporated into the *E. coli* chromosome via integration into the bacterial *attB* site. Such an event was dependent on the expression of a phage Integrase ^1^ Thus, genes expressed from endogenous or regulatable promoters could be delivered into the stable confines of the chromosome in single copy, without the worry of the physiological artifacts of excessive gene copy number or the instability of self-autonomous replication vectors. Site-specific recombination (SSR) systems were the basis for developing the Lambda InCh method for transferring exogenous genes to the *E. coli attB* site ^2^, the series of CRIM plasmids that take advantage of a variety of different phage *attB* sites in *E. coli*^3^, and many technical modifications of these SSR systems and their substrates for use in a variety of metabolic and genetic engineering protocols in a variety of organisms ^4-8^.

In addition to these site-specific recombination systems, general homologous recombination (HR) systems of phage, such as the Red system of phage lambda, and the RecET systems of Rac prophage and MTb phage Che9c, have been widely used for genetic engineering in a variety of bacteria ^9-15^; for reviews, see ^16-18^. Termed “recombineering”, these procedures have used both dsDNA PCR substrates and ssDNA oligos for generating bacterial chromosomal modifications in *E. coli*, and in various bacterial pathogens including enterohemorrhagic *E. coli., Salmonella enterica, Shigella flexneri, Pseudomonas aeruginosa, Yersinia pestis*, and *M. tuberculosis* ^14, 15, 19-26^. These general phage recombination systems have also been used for the development of new methodologies for metabolic engineering of industrial microorganisms ^27-36^. The Red system annealase λ Beta protein, in particular, has been employed in many protocols for the systematic modification of *E. coli*, for instance, by electroporation of oligos targeting the promoter regions of genomic targets allowing for accelerated evolution of *E. coli* for specific metabolic engineering purposes ^37^. In concert with these homologous recombination systems, SSR systems described above have been employed to remove selectable drug cassettes for the construction of marker-less gene deletions and fusions ^10, 12, 38^

The new system described in this report couples the recombinogenic annealing of an oligo and the site-specific insertion of a non-replicating plasmid into a one-step procedure for generating chromosomally tagged genes, deletions, or promoter replacements in *M. smegmatis* or *M. tuberculosis* (MTb). It is the first general chromosomal engineering technique that produces a drug-selectable recombinant that does not require the use of either target-specific dsDNA plasmids or PCR-generated recombination substrates. The only target-specific substrate requirement for gene modification is a chemically synthesized oligo. This “targeting oligo” carries the ssDNA version of the Bxb1 phage *attP* site (48 bases) flanked by 45-70 bases of homology to the chromosomal target, and is co-electroporated with a non-replicating “payload plasmid” that contains a Bxb1 *attB* site. The host (*M. smegmatis* or *M. tuberculosis*) contains a plasmid that expresses both the Che9c phage RecT annealase and the Bxb1 phage Integrase. RecT promotes annealing of the targeting oligo to the lagging strand template of the replication fork, thus placing the *attP* site into a precise location in the chromosome dictated by the oligo sequence. *In the same outgrowth period*, Bxb1 integrase promotes site-specific recombination between the co-electroporated attB-containing payload plasmid and the oligo-derived *attP* site. In this system, the sequence of the oligo defines the position of the insertion site and the plasmid delivers the payload. For knockouts, the oligo is designed so that *attP* replaces the target gene. For C-terminal tags, the oligo is designed to insert *attP* at the end of the coding sequence. In this case, the type of C-terminal tag desired is defined by the selection of the plasmid to be co-electroporated with the oligo from a library of pre-existing payload plasmids. Inherent in this system, an oligo designed to create a C-terminal tag can be used with multiple plasmids to create fluorescent, degradation, and epitope tagged fusions. This new gene modification scheme is called ORBIT (for <u>O</u>ligo <u>R</u>ecombineering followed by <u>B</u>xb1-<u>I</u>ntegrase <u>T</u>argeting). We describe the use of the system to generate over 100 gene knockouts and fusions in *M. smegmatis* and *M. tuberculosis* at high efficiency.

## RESULTS

### RecT-promoted Oligo-mediated Recombineering – 60 bp insertion

Two different types of recombineering methodologies have been applied to mycobacteria, but neither represents a broadly generalizable approach for genome engineering. Target-specific dsDNA substrates can be used to make diverse and selectable mutations. However, even the smallest useful recombination substrates consist of both a selectable marker and ~500 bp of flanking homology to the chromosome. These dsDNA constructs are cumbersome to generate for each desired mutation, and even these large substrates recombine at relatively low-efficiency. In contrast, single-stranded oligos are easily synthesized and can be used to alter one or a few bases of the chromosome at high efficiency. However, these mutations are generally not selectable and therefore difficult to isolate. An ideal method would leverage easily synthesized and highly efficient oligonucleotide substrates to make selectable mutations. We sought to accomplish this by encoding a phage attachment site (*attP*) in the oligo that could be used to integrate a selectable marker at a specific chromosomal site via site-specific recombination.

To test this idea, an assay was designed to measure the frequency of incorporation of an oligo containing an insertion of approximately the size of a 48 bp *attP* site into the mycobacterial chromosome. For this purpose, a hygromycin resistance gene, with an internal 60 bp deletion, was integrated into the L5 phage attachment site of the *M. smegmatis* chromosome. Oligos (180-mers) targeting the lagging strand template of the impaired Hyg^R^ marker and containing the missing 60 bases (as well as 60 bases flanking the deletion site) were electroporated into *M. smegmatis* expressing the Che9c RecT annealase from the anhydrotetracycline (ATc)-inducible Ptet promoter (pKM402) ^39^. The frequency of oligo incorporation was determined as the number of Hyg^R^ transformants among the survivors of electroporation. The number of Hyg^R^ transformants generated varied from less than 10 to over 300, depending on the amount of oligo used (Fig. 1b). At about 1 μg of oligo, the total number of Hyg^R^ recombinants plateaued at approximately 350 transformants per electroporation, which corresponds to a frequency of 2 x 10^-6^ recombinants per survivor of electroporation. In an experiment where the target contained a 1 bp change creating a premature stop codon in the *hyg*-resistance gene, a 60 base oligo was used to restore Hyg resistance at a frequency of 4 x 10^-4^ recombinants per survivor of electroporation. Thus, the RecT annealase is capable of integrating an oligo that contains a 60 base insertion into the mycobacterial chromosome, albeit at a frequency which is ~500-fold lower relative to a single base pair change.

**Figure 1.**
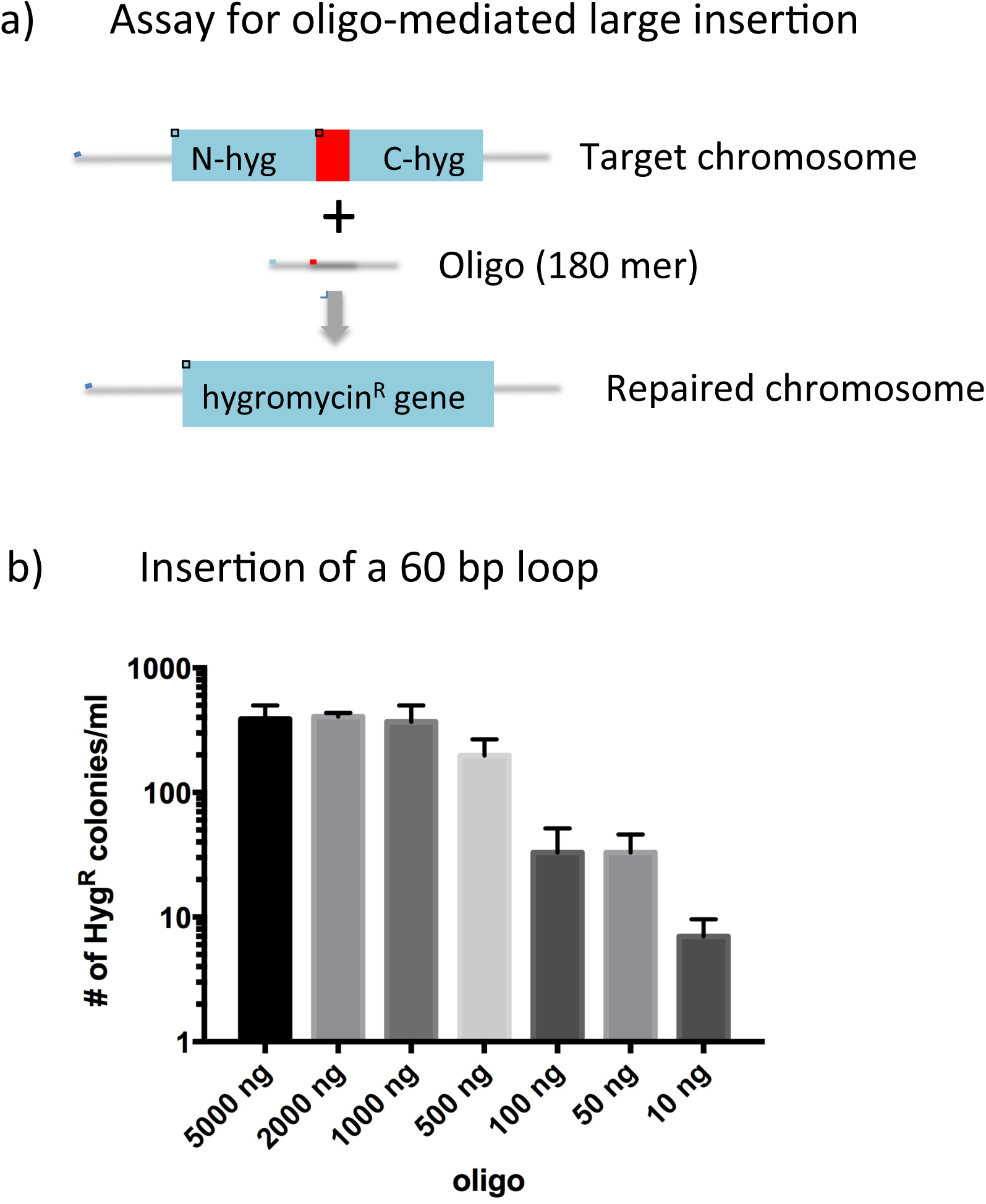
RecT-promoted oligo-mediated 60 base insertion. (a) Diagram of oligo-mediated recombineering of a chromosomal target in *M. smegmatis*. An integrating plasmid (pKM433) at the L5 phage attachment site contains a mutated *hyg*-resistance gene due to an internal 60 bp deletion (red square). Electroporation of an oligo containing the 60 bases missing in the target gene, along with 60 bp of flanking DNA on each side, is electroporated into cells expressing the Che9c RecT function from pKM402. (b) After induction of RecT and preparing the cells for transformation (as described in the Methods section), the cells were electroporated with various amounts of an oligo (180 mer) that spans the 60 bp deletion of the Hyg-resistance cassette in pKM433. Cells were grown out overnight and 0.5 ml was plated on LB-Hyg plates. The experiment was performed in triplicate; standard errors are shown.

### Development of ORBIT

To convert the recombineering of an oligo into a selectable event, we determined if co-electroporation of the *attP*-containing oligo with an *attB*-containing non-replicating vector (Hyg^R^) into a cell that expresses both the RecT annealase and the phage Bxb1 Integrase would allow for both homologous and site-specific recombination events to occur within the same outgrowth period. Since the oligo is designed to direct the integration of the genetic information contained in the non-replicating plasmid, these elements were termed, “targeting oligo” and “payload plasmid”.

Two plasmids were generated to test this methodology (Fig. 2). pKM444 produces the recombination functions. This mycobacterial shuttle vector expresses both the Che9c phage RecT annealase and the Bxb1 phage integrase (Int) from the Ptet promoter (Fig. 2a). pKM446 is a payload plasmid that will not replicate in mycobacteria. This vector encodes a *hyg* resistance marker for selection in mycobacteria and a Bxb1 *attB* site (Fig. 2b). Adjacent to the *attB* site is a sequence encoding both a Flag tag and a DAS+4 peptide tag designed to be in frame with a targeted chromosomal gene following integration of the plasmid. The DAS+4 tag directs a fusion protein for degradation via the ClpXP system upon expression of the SspB adapter protein ^40, 41^. Targeting oligos were designed to direct the integration of *attP* to the 3’ ends of the *M. smegmatis recA*, *DivIVA* and *leuB* genes, just in front of the stop codon. A site-specific recombination event between the inserted *attP* and the coelectroporated pKM446 would then generate a DAS+4 fusion to these target genes (see Supplementary Table 1 for the list of oligos used in this study). Targeting oligos, which anneal to the lagging strand template of the replication fork, were co-electroporated with the pKM446 payload plasmid into *M. smegmatis* that expressed both Che9c RecT annealase and Bxb1 Integrase. Among the Hyg^R^ colonies resulting from these transformations, 9 out of 12 candidates tested by PCR contained the expected recombination structure, in which pKM446 was inserted between *attR* and *attL* sites at the predicted oligo-directed integration site (Fig. 2 c-e). The fusion of the target genes to the Flag-DAS+4 degradation tag was verified by sequencing the PCR products of the 5’ junctions. The sequence of the recA-Flag-DAS+4 fusion generated by ORBIT is shown in Supplementary Fig. 1.

**Figure 2.**
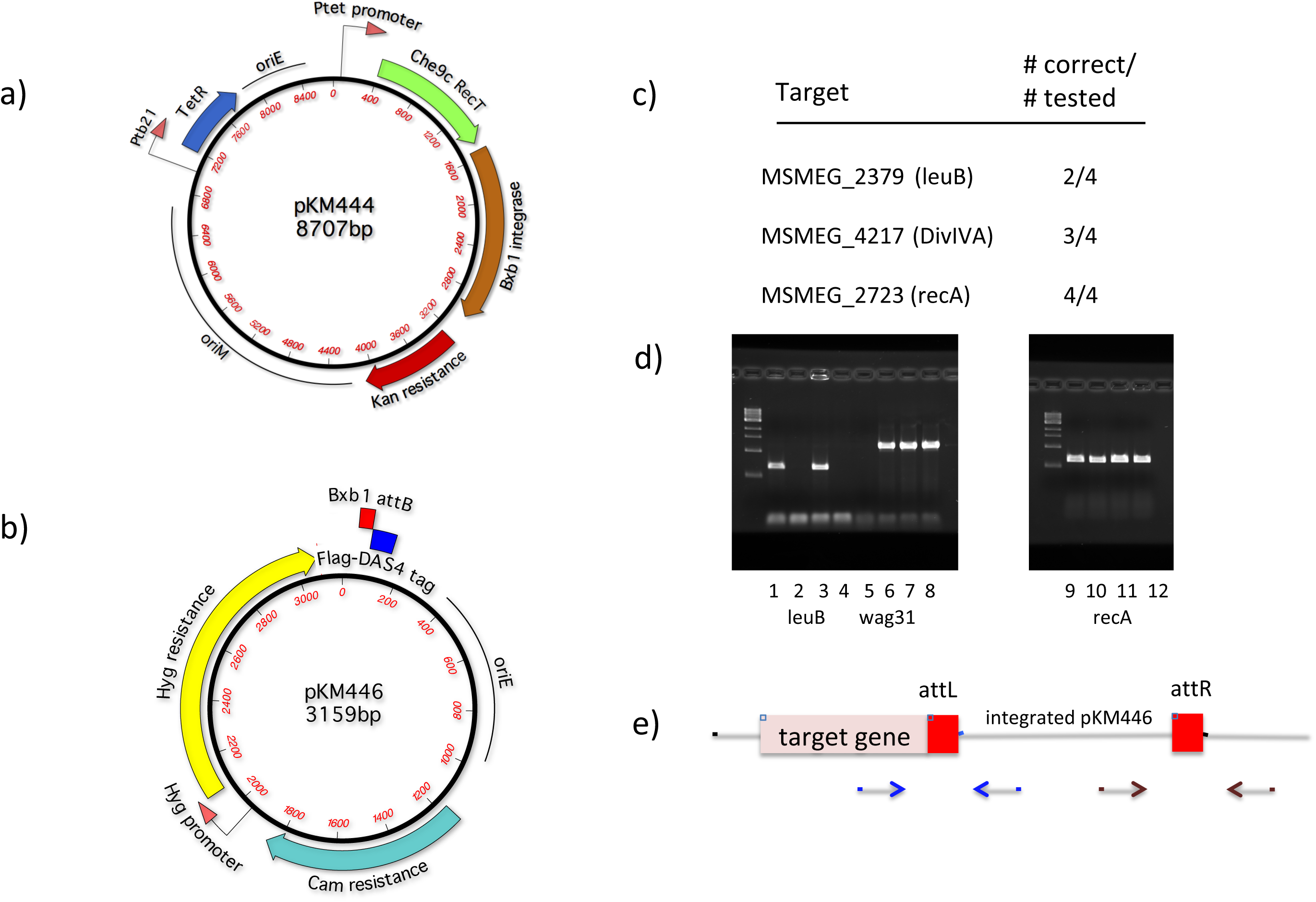
Plasmids constructed for ORBIT. (a) Construct pKM444 expresses the Che9c phage RecT annealase and the Bxb1 phage Integrase, both driven from the Ptet promoter. A similar construct (pKM461) contains (in addition) the *sacRB* genes for curing the plasmid following gene modification. (b) One of the ORBIT payload plasmids (pKM446) used for integration into the chromosomal *attP* site created by an oligo. In this case, the plasmid payload contains a Flag-DAS+4 degradation tag that will be fused to the 3’ end of the target gene. (c) List of 3 genes in *M. smegmatis* targeted for C-terminal tagging. Following the ORBIT protocol for each target gene, total numbers of colonies obtained (from multiple trials) ranged between 10-100 CFU/ml. Electroporations with payload plasmid only (no targeting oligo) gave, on average, 5-fold fewer total numbers of colonies. The number of correct recombinants (out of 4 candidates tested) for each target gene is shown. (d) PCR analysis of the 5’ junctions of each candidate tested. (e) Primer positions for verification by PCR of the recombinants are shown; 5’ junction (blue arrows), 3’ junction (brown arrows). In each case where a 5’ junction was verified, the 3’ junction was also verified (not shown). The 5’ junctions were confirmed by DNA sequencing.

To test the functionality of the tag and verify that the mutated locus was the only copy of the targeted gene in the cell, we took advantage of the protein degradation system for controlling the expression of DAS+4-tagged proteins in *M. smegmatis* and *M. tuberculosis* ^42-44^ In this system, the degradation of the DAS+4 tagged protein is promoted by expression of the *E. coli* SspB protein. Two different *sspB* expression systems were used, which induce degradation of the target protein either by adding ATc (Tet-OFF) or removing ATc (Tet-ON).

Cells containing the *recA*-DAS+4 fusion were transformed with pGMCgS-TetON-18, allowing the induction of SspB in the absence of ATc; RecA function was quantified using a UV resistance assay. When grown in the presence of ATc, the *recA*-DAS+4-tagged strain containing either the SspB-expressing plasmid or control plasmid showed similar levels of UV sensitivity (Fig. 3a right). In contrast, the sensitivity of the tagged strain with SspB was enhanced in the absence of ATc (Fig. 3a left). The *divIVA* and *leuB* DAS+4-tagged strains were transformed with pGMCgS-TetOFF-18, which produces SspB upon ATc addition. In both cases, addition of ATc inhibited the growth of these mutants, consistent with the essentiality of these genes on the media used in this study (Fig. 3b & c). The defect of LeuB depletion could be reversed by the addition of leucine to the plates, verifying that that this phenotype was linked to the engineered mutation (not shown). Thus, the ORBIT method could generate function-altering mutations without the need to construct either target-specific plasmids or long dsDNA recombineering substrates.

**Figure 3.**
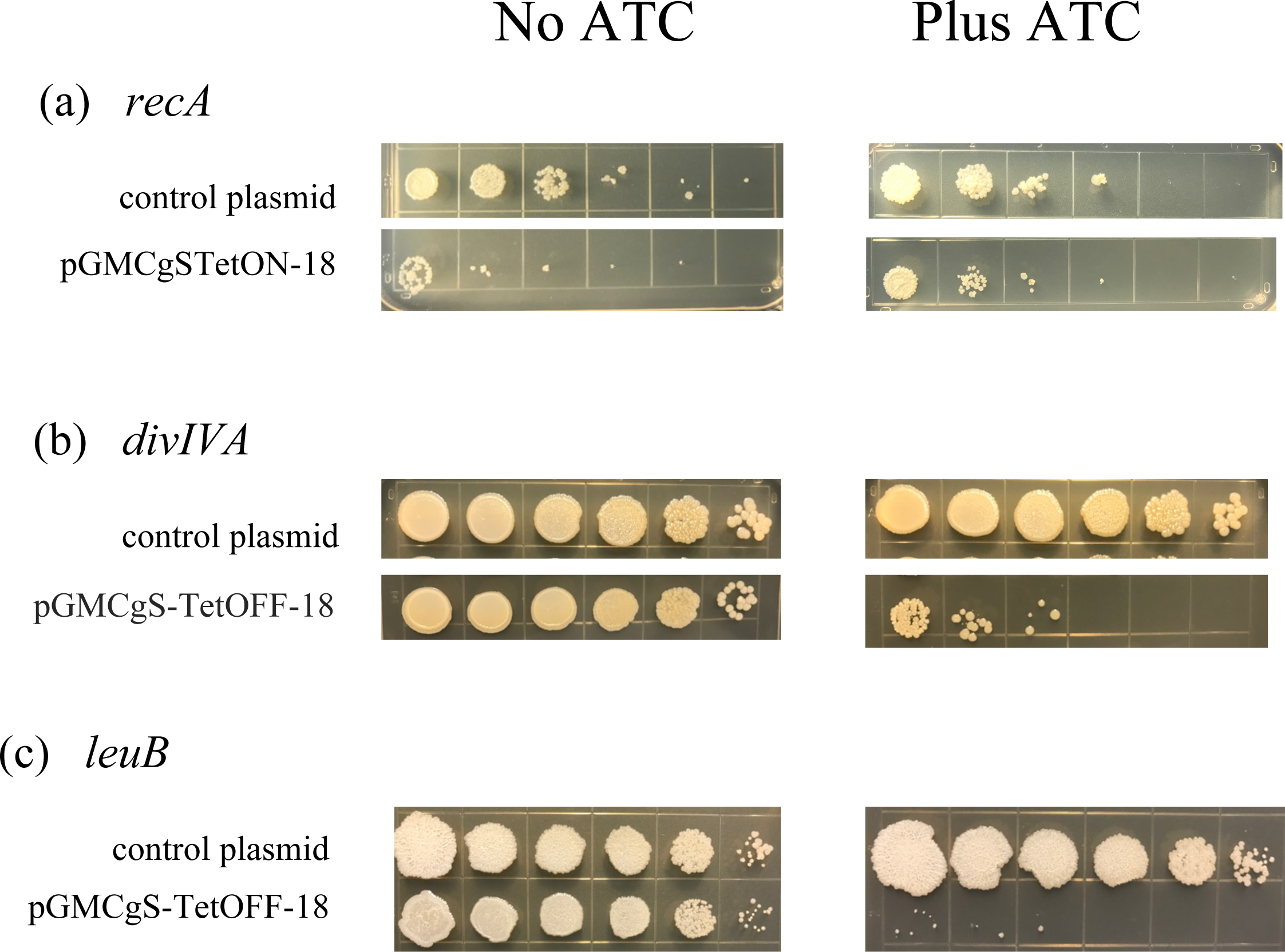
Knockdown phenotypes of ORBIT-generated DAS+4-tagged strains. The growth phenotypes of the Flag-DAS+4 tagged strains constructed above were analyzed after transformation of an SspB-expressing plasmid. **(a)** The *recA*-Flag-DAS+4 strain was transformed with an SspB-producing plasmid pGMCgS-TetON-18 (strep^R^) under control of the reverse TetR repressor. In this scenario, RecA is proteolyzed in the *absence* of anhydrotetracycline (ATc). Ten-fold serial dilutions of the cells were spotted on LB-strep plates and the cells were exposed to 20 J/m2 of UV. In the absence of ATc, increased sensitivity of *recA*-Flag-DAS+4 strain containing pGMCgTetON-18 (relative to the tagged strain containing a control plasmid) is observed (left). In the presence of ATc, both strains show similar UV sensitivities (right). **(b)** The *DivIVA*-Flag-DAS+4 strain was transformed with an SspB-producing plasmid pGMCgS-TetOFF-18 (strep^R^) under control of the wild type TetR repressor. In this case, DivIVA is expected to be depleted in the presence of ATc. In the absence of ATc, both cultures grow well on LB-strep plates. In the presence of ATc, growth sensitivity is observed for the *DivIVA*-Flag-DAS+4 strain containing the SspB-producer. **(c)** Same as B, except that *leuB* is the target and the cells are plated on 7H9 plates.

The scheme of using both RecT and Bxb1 Integrase simultaneously to promote modification of a chromosomal target gene is diagramed in Fig. 4; the process is called ORBIT (for <u>O</u>ligo-mediated <u>R</u>ecombineering followed by <u>B</u>xb1 <u>I</u>ntegrase <u>T</u>argeting). In theory, targeting oligos could also be designed to delete a portion of the chromosome during the ORBIT reaction (Fig. 4, right side) to create deletions. Demonstrations of such ORBIT-promoted knockouts are described bellow.

**Figure 4.**
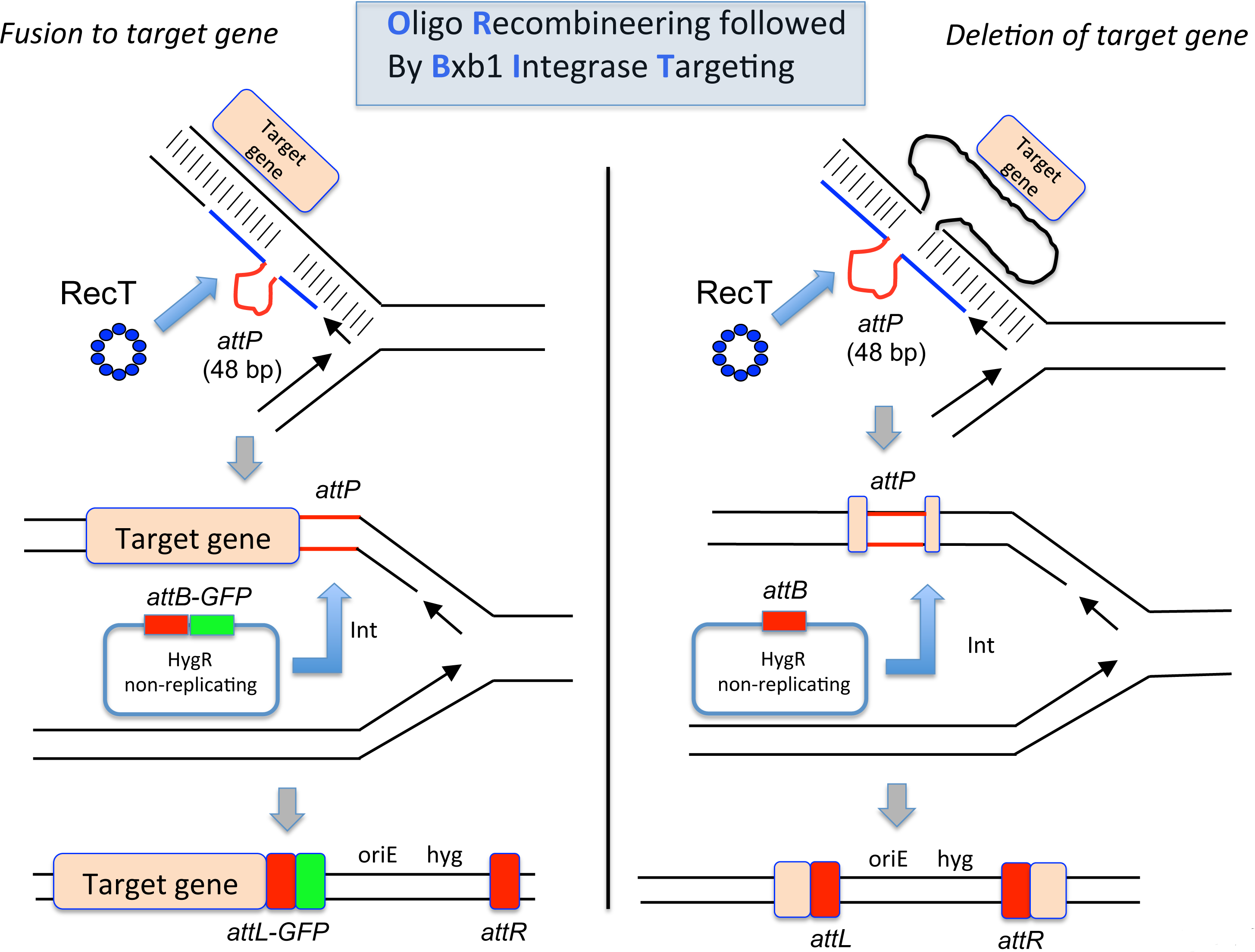
ORBIT-promoted gene alteration. The site of action occurs at the replication fork. An oligomer containing a single-stranded version of the Bxb1 *attP* site (top pictures, red lines) is co-electroporated with an *attB*-containing non-replicating plasmid into a mycobacterial host cell expressing both RecT annealase and Bxb1 Integrase. RecT promotes annealing of the oligo to the lagging strand template. Following DNA replication through this region, an *attP* site is formed in the chromosome (middle pictures). In the same outgrowth period, Bxb1 Integrase promotes site-specific insertion of the plasmid into the chromosome (*attB* x *affP*). Left side: The oligo is designed so that *attP* is inserted just before the stop codon. The integration event fuses the GFP tag in-frame to the 3’ end of the target gene (with an *attL* site in-frame between them); the recombinant is selected for by Hyg^R^. Right side: The oligo is designed so that *attP* replaces the target gene and the plasmid integration event allows hygromycin resistance to be used to select for the knockout.

### Parameters of ORBIT-promoted gene targeting

Two features of ORBIT that were optimized are the length of the homologous arms (HAs) in the targeting oligo, and the relative amounts of oligo and non-replicating plasmid used for coelectroporation. The HAs of a *polA*-targeting oligo were varied from 50 to 70 bases (oligo lengths, including *attP*, were from 148 to 188 bases). The oligos were mixed with 200 ng of pKM446 and transformed into *M. smegmatis* containing pKM444. While there is variability in the number of Hyg-resistant transformants from each electroporation, oligos containing longer HAs produced more transformants (Fig. 5a). Below 40 base pair HAs, no Hyg^R^ transformants are observed above the number seen in the no oligo control.

**Figure 5.**
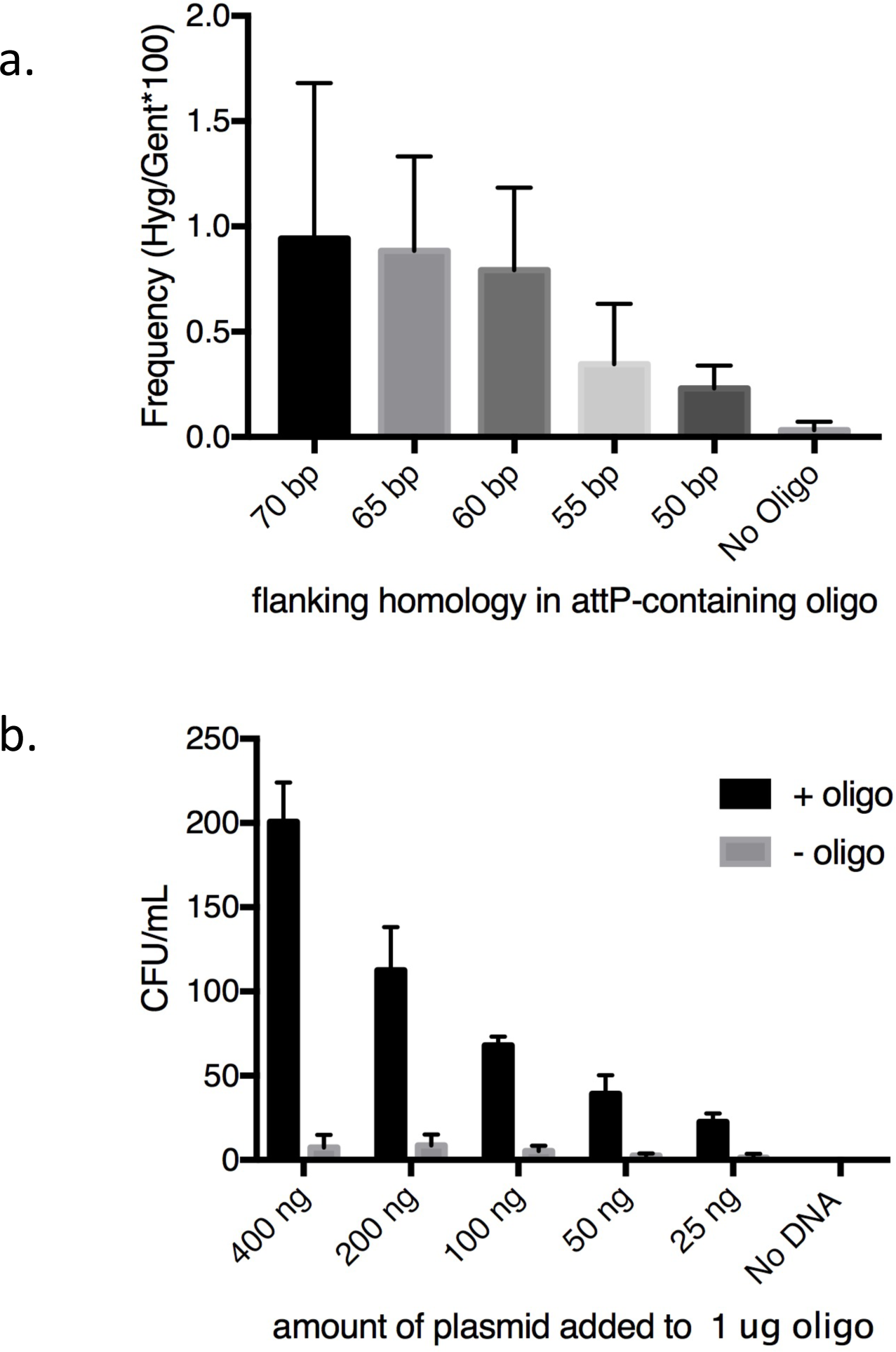
Parameters of the ORBIT process. **A.** The amount of target homology flanking the *attP* site in an oligo designed to create a *polA*-Flag-DAS+4 fusion in *M. smegmatis* was examined as a function of recombinant formation (Hyg^R^). One microgram of each oligo was electroporated with 200 ng of pKM446. The frequency of targeting is expressed as the percentage of Hyg^R^ transformants following integration of pKM446 relative to a transformation control (20 ng of Gen^R^ plasmid pKM390). Experiments were performed in triplicate; standard errors are shown. **B.** Colony counts were measured after electroporation of 1 μg of an oligo with 70 base flanks (designed to create a *polA*-Flag-DAS+4 fusion) with various amounts of pKM446. CFU/ml was measured following overnight growth of the electroporation mixtures in 2ml LB. Experiments were performed in triplicate; standard errors are shown.

We also examined the optimal ratio of oligo to plasmid. An optimal concentration of targeting oligo (1 μg) was co-electroporated with various amounts of the payload plasmid pKM446. More Hyg^R^ transformants are observed when more plasmid is used, but there is also a small increase in the number of oligo-independent transformants that presumably represent illegitimate recombinants (Fig. 5b). PCR screening confirmed that 39/40 Hyg-resistant transformants recovered in this experiment (using 8 candidates of each transformation), represented the desired oligo-directed recombination events. As a rule, we generally combined 1 μg of oligo with 200 ng of plasmid in the transformations described below.

### ORBIT-promoted knockdowns and knockouts in *M. smegmatis and M. tuberculosis*

To determine if ORBIT-promoted modifications could be generally engineered throughout the chromosome, we targeted a variety of genes in *M. tuberculosis* and *M. smegmatis*. For C-terminal tags, the *attP* site was placed just in front of the stop codon of the target gene. For knockouts, the *attP* site was flanked by 60-70 bases, which included the first and last 10 codons of the target gene (including the start and termination codons), which is expected to result in the deletion of intervening chromosomal sequence. Overall, we have made over 100 strains where the target genes have either been deleted or C-terminally tagged (see Fig. 6a & b and Tables 1 & 2). For most of these targets, 5 to 50 colonies were typically observed after plating 0.5 ml of the overnight outgrowth. Usually, only 2-4 Hyg^R^ candidates needed to be analyzed by PCR to identify at least one strain that contained the payload plasmid in the site designated by the targeting oligo. Most of the targeting oligos used for these genomic modifications contained either 60 or 70 bases of flanking homology. An oligo targeting the *aceE* gene containing only 45 bases homology on each side of *attP* also produced the desired recombinant in 4 of 6 clones. However, lowering flanking homologies below 40 bases decreased the percentage of correct recombinants dramatically, largely as a result of an increased number of illegitimate recombination events.

**Figure 6.**
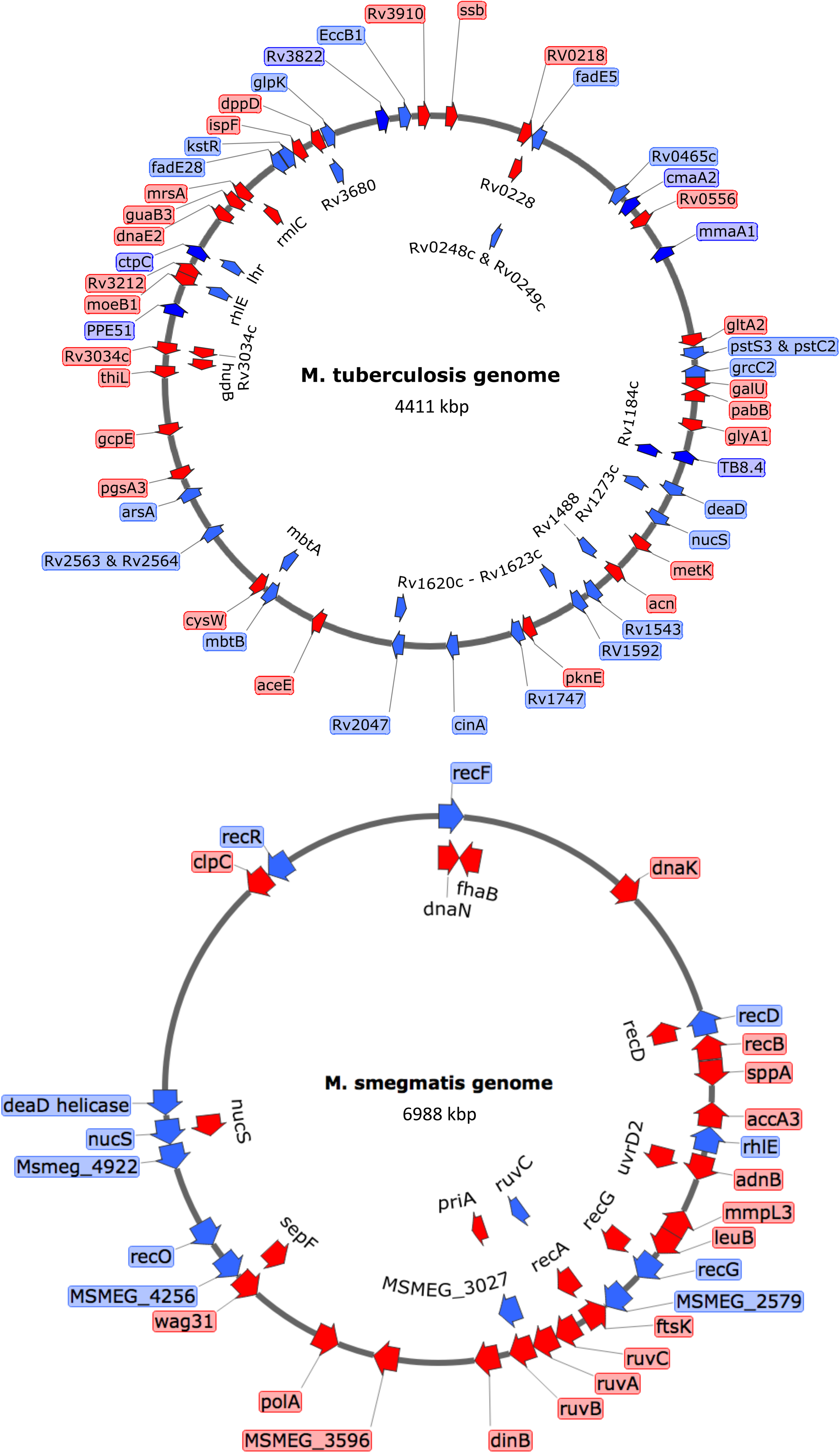
ORBIT-generated insertions and deletions in *M. tuberculosis* and *M. smegmatis*. Target gene deletions and tags were placed at a variety of positions in the chromosomes of both *M. tuberculosis* (a) and *M. smegmatis* (b). In most cases, the oligos contained an *attP* site flanked by 70 bases of target homology. Insertions (either DAS+4, GFP or His-Flag) are shown in red and deletions are shown in blue. A description of all the types of modifications performed by ORBIT are shown in Table 1 (for *M. smegmatis*) and Table 2 (for *M. tuberculosis*).

**Table 1.**
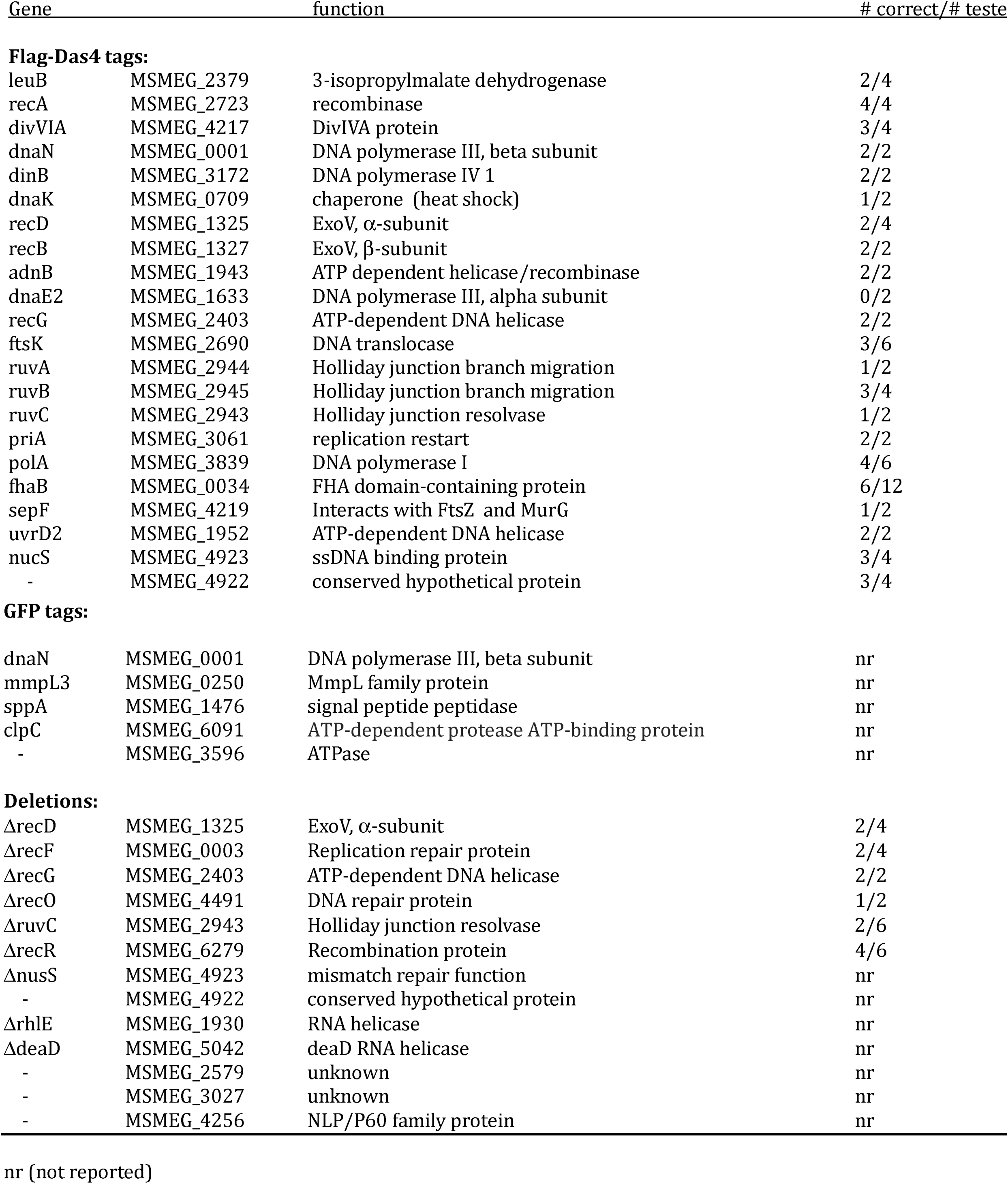
ORBIT-promoted *M. smegmatis* modifications.

| Gene |  | function | # correct/# tested |
| --- | --- | --- | --- |
| Flag-Das4 tags: |  |  |  |
| leuB | MSMEG_2379 | 3-isopropylmalate dehydrogenase | 2/4 |
| recA | MSMEG_2723 | recombinase | 4/4 |
| divVIA | MSMEG_4217 | DivIVA protein | 3/4 |
| dnaN | MSMEG_0001 | DNA polymerase III, beta subunit | 2/2 |
| dinB | MSMEG_3172 | DNA polymerase IV 1 | 2/2 |
| dnaK | MSMEG_0709 | chaperone (heat shock) | 1/2 |
| recD | MSMEG_1325 | ExoV, α-subunit | 2/4 |
| recB | MSMEG_1327 | ExoV, β-subunit | 2/2 |
| adnB | MSMEG_1943 | ATP dependent helicase/recombinase | 2/2 |
| dnaE2 | MSMEG_1633 | DNA polymerase III, alpha subunit | 0/2 |
| recG | MSMEG_2403 | ATP-dependent DNA helicase | 2/2 |
| ftsK | MSMEG_2690 | DNA translocase | 3/6 |
| ruvA | MSMEG_2944 | Holliday junction branch migration | 1/2 |
| ruvB | MSMEG_2945 | Holliday junction branch migration | 3/4 |
| ruvC | MSMEG_2943 | Holliday junction resolvase | 1/2 |
| priA | MSMEG_3061 | replication restart | 2/2 |
| polA | MSMEG_3839 | DNA polymerase I | 4/6 |
| fhaB | MSMEG_0034 | FHA domain-containing protein | 6/12 |
| sepF | MSMEG_4219 | Interacts with FtsZ and MurG | 1/2 |
| uvrD2 | MSMEG_1952 | ATP-dependent DNA helicase | 2/2 |
| nucS | MSMEG_4923 | ssDNA binding protein | 3/4 |
| - | MSMEG_4922 | conserved hypothetical protein | 3/4 |
| **GFP tags:** |  |  |  |
| dnaN | MSMEG_0001 | DNA polymerase III, beta subunit | nr |
| mmpL3 | MSMEG_0250 | MmpL family protein | nr |
| sppA | MSMEG_1476 | signal peptide peptidase | nr |
| clpC | MSMEG_6091 | ATP-dependent protease ATP-binding protein | nr |
| - | MSMEG_3596 | ATPase | nr |
| **Deletions:** |  |  |  |
| ΔrecD | MSMEG_1325 | ExoV, α-subunit | 2/4 |
| ΔrecF | MSMEG_0003 | Replication repair protein | 2/4 |
| ΔrecG | MSMEG_2403 | ATP-dependent DNA helicase | 2/2 |
| ΔrecO | MSMEG_4491 | DNA repair protein | 1/2 |
| ΔruvC | MSMEG_2943 | Holliday junction resolvase | 2/6 |
| ΔrecR | MSMEG_6279 | Recombination protein | 4/6 |
| ΔnusS | MSMEG_4923 | mismatch repair function | nr |
| - | MSMEG_4922 | conserved hypothetical protein | nr |
| ΔrhIE | MSMEG_1930 | RNA helicase | nr |
| ΔdeaD | MSMEG_5042 | deaD RNA helicase | nr |
| - | MSMEG_2579 | unknown | nr |
| - | MSMEG_3027 | unknown | nr |
| - | MSMEG_4256 | NLP/P60 family protein | nr |

nr (not reported)

**Table 2.**
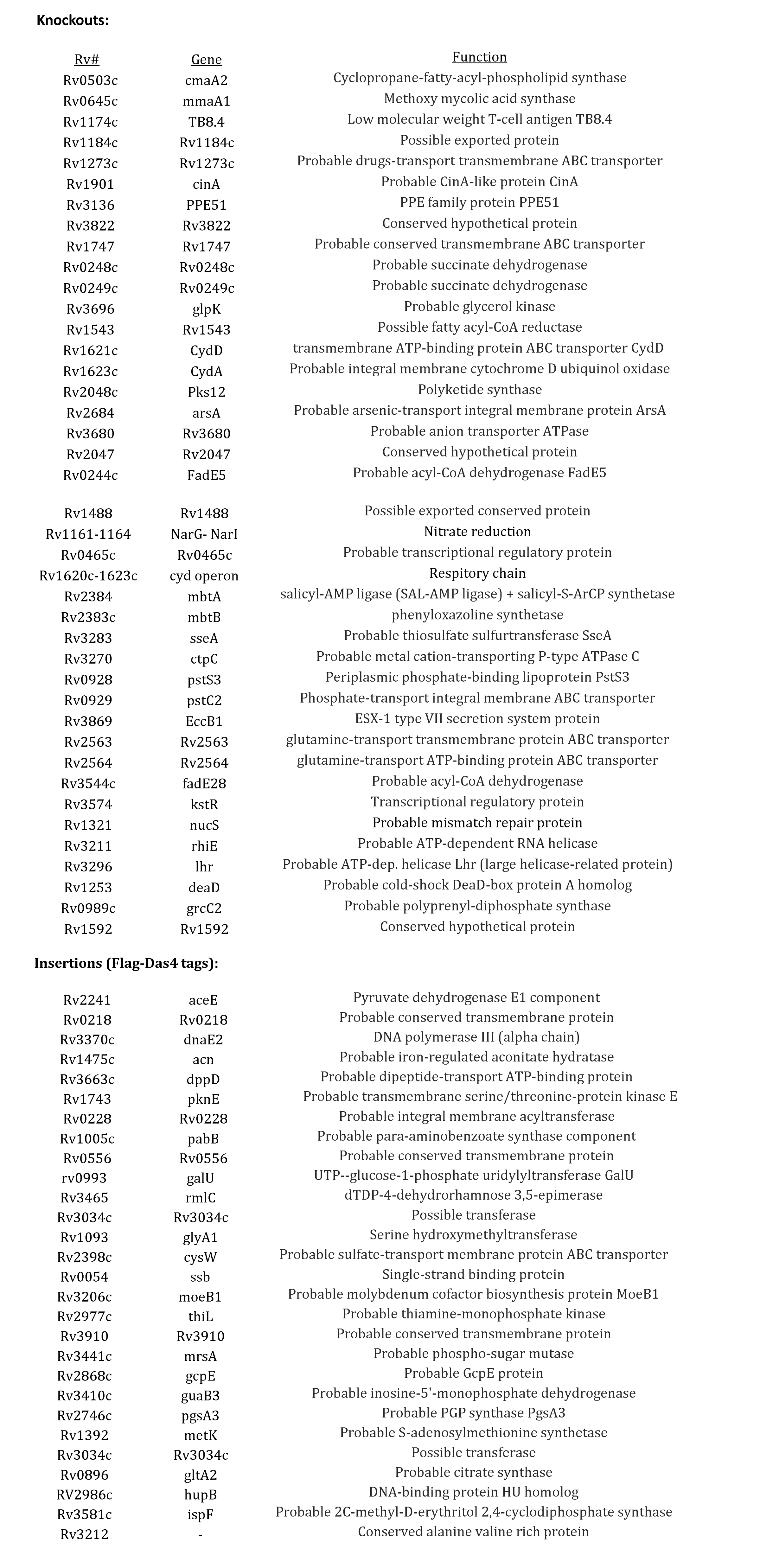
ORBIT-promoted *M. tuberculosis* modifications.

| <u>Rv#</u> | <u>Gene</u> | <u>Function</u> |
| --- | --- | --- |
| Rv0503c | cmaA2 | Cyclopropane-fatty-acyl-phospholipid synthase |
| Rv0645c | mmaA1 | Methoxy mycolic acid synthase |
| Rv1174c | TB8.4 | Low molecular weight T-cell antigen TB8.4 |
| Rv1184c | Rv1184c | Possible exported protein |
| Rv1273c | Rv1273c | Probable drugs-transport transmembrane ABC transporter |
| Rv1901 | cinA | Probable CinA-like protein CinA |
| Rv3136 | PPE51 | PPE family protein PPE51 |
| Rv3822 | Rv3822 | Conserved hypothetical protein |
| Rv1747 | Rv1747 | Probable conserved transmembrane ABC transporter |
| Rv0248c | Rv0248c | Probable succinate dehydrogenase |
| Rv0249c | Rv0249c | Probable succinate dehydrogenase |
| Rv3696 | glpK | Probable glycerol kinase |
| Rv1543 | Rv1543 | Possible fatty acyl-CoA reductase |
| Rv1621c | CydD | transmembrane ATP-binding protein ABC transporter CydD |
| Rv1623c | CydA | Probable integral membrane cytochrome D ubiquinol oxidase |
| Rv2048c | Pks12 | Polyketide synthase |
| Rv2684 | arsA | Probable arsenic-transport integral membrane protein ArsA |
| Rv3680 | Rv3680 | Probable anion transporter ATPase |
| Rv2047 | Rv2047 | Conserved hypothetical protein |
| Rv0244c | FadE5 | Probable acyl-CoA dehydrogenase FadE5 |
| Rv1488 | Rv1488 | Possible exported conserved protein |
| Rv1161-1164 | NarG- NarI | Nitrate reduction |
| Rv0465c | Rv0465c | Probable transcriptional regulatory protein |
| Rv1620c-1623c | cyd operon | Respiratory chain |
| Rv2384 | mbtA | salicyl-AMP ligase (SAL-AMP ligase) + salicyl-S-ArCP synthetase |
| Rv2383c | mbtB | phenyloxazoline synthetase |
| Rv3283 | sseA | Probable thiosulfate sulfurtransferase SseA |
| Rv3270 | ctpC | Probable metal cation-transporting P-type ATPase C |
| Rv0928 | pstS3 | Periplasmic phosphate-binding lipoprotein PstS3 |
| Rv0929 | pstC2 | Phosphate-transport integral membrane ABC transporter |
| Rv3869 | EccB1 | ESX-1 type VII secretion system protein |
| Rv2563 | Rv2563 | glutamine-transport transmembrane protein ABC transporter |
| Rv2564 | Rv2564 | glutamine-transport ATP-binding protein ABC transporter |
| Rv3544c | fadE28 | Probable acyl-CoA dehydrogenase |
| Rv3574 | kstR | Transcriptional regulatory protein |
| Rv1321 | nucS | Probable mismatch repair protein |
| Rv3211 | rhiE | Probable ATP-dependent RNA helicase |
| Rv3296 | lhr | Probable ATP-dep. helicase Lhr (large helicase-related protein) |
| Rv1253 | deaD | Probable cold-shock DeaD-box protein A homolog |
| Rv0989c | grcC2 | Probable polyprenyl-diphosphate synthase |
| Rv1592 | Rv1592 | Conserved hypothetical protein |
| Insertions (Flag-Das4 tags): |  |  |
| Rv2241 | aceE | Pyruvate dehydrogenase E1 component |
| Rv0218 | Rv0218 | Probable conserved transmembrane protein |
| Rv3370c | dnaE2 | DNA polymerase III (alpha chain) |
| Rv1475c | acn | Probable iron-regulated aconitate hydratase |
| Rv3663c | dppD | Probable dipeptide-transport ATP-binding protein |
| Rv1743 | pknE | Probable transmembrane serine/threonine-protein kinase E |
| Rv0228 | Rv0228 | Probable integral membrane acyltransferase |
| Rv1005c | pabB | Probable para-aminobenzoate synthase component |
| Rv0556 | Rv0556 | Probable conserved transmembrane protein |
| rv0993 | galU | UTP--glucose-1-phosphate uridylyltransferase GalU |
| Rv3465 | rmlC | dTDP-4-dehydrorhamnose 3,5-epimerase |
| Rv3034c | Rv3034c | Possible transferase |
| Rv1093 | glyA1 | Serine hydroxymethyltransferase |
| Rv2398c | cysW | Probable sulfate-transport membrane protein ABC transporter |
| Rv0054 | ssb | Single-strand binding protein |
| Rv3206c | moeB1 | Probable molybdenum cofactor biosynthesis protein MoeB1 |
| Rv2977c | thiL | Probable thiamine-monophosphate kinase |
| Rv3910 | Rv3910 | Probable conserved transmembrane protein |
| Rv3441c | mrsA | Probable phospho-sugar mutase |
| Rv2868c | gcpE | Probable GcpE protein |
| Rv3410c | guaB3 | Probable inosine-5'-monophosphate dehydrogenase |
| Rv2746c | pgsA3 | Probable PGP synthase PgsA3 |
| Rv1392 | metK | Probable S-adenosylmethionine synthetase |
| Rv3034c | Rv3034c | Possible transferase |
| Rv0896 | gltA2 | Probable citrate synthase |
| RV2986c | hupB | DNA-binding protein HU homolog |
| Rv3581c | ispF | Probable 2C-methyl-D-erythritol 2,4-cyclodiphosphate synthase |
| Rv3212 | - | Conserved alanine valine rich protein |

In order to cure recombinants of the RecT-Int producing plasmid following modification, a *sacB*-containing derivative of pKM444 was constructed (pKM461). A number of genes were tagged in a pKM461-bearing strain (including *recG*, *dnaN* and *ftsK*), then plated directly on Hyg-sucrose plates to both select for the recombinant and to cure the strain of the RecT-Int producer. This process appears to produce fewer colonies than previously observed with pKM444, but can rapidly produce plasmid-free mutants.

Genetic modifications with ORBIT should be very stable. An excision event promoted by the Bxb1 SSR system requires an additional factor called gp47 ^45^, which is not present in the hosts used for these experiments. The stability of ORBIT-mediated modifications was verified by observing that the integrated payload plasmid, in a number of *M. smegmatis* mutants generated in this study, was not lost after more than 100 generations of growth in the absence of drug selection. The one exception was *dnaK*-Flag-DAS+4, a fusion that could be constructed, but was lost spontaneously in subsequent outgrowth periods, as evidenced by the loss of plasmid-chromosome junctions. Apparently, this important chaperone cannot sustain the presence of the tag on its C-terminal end, especially during a stress period such as selection on sucrose for the curing of *sacB*-containing plasmids.

### Expanding the library of ORBIT-mediated modifications

In order to expand the types of modifications that can be made via ORBIT, we built a set of payload plasmids all containing the Bxb1 *attB* site fused to different types of tags. In addition to the Flag-DAS+4 plasmids, we generated plasmids to create C-terminal targeted fusions with combinations of eGFP, mVenus, SNAP, CLIP, Myc and His epitopes, and TEV cleavage sites. These tags facilitate both protein localization by fluorescence and tandem affinity purification (Table 3). In addition, we created plasmids to generate chromosomal knockouts and promoter replacements using either hygromycin or zeocin resistance as a selection for the recombinant (see Table 3). Any number of payload plasmids can be matched with a single targeting oligo to generate a variety of functional gene modifications.

**Table 3.**
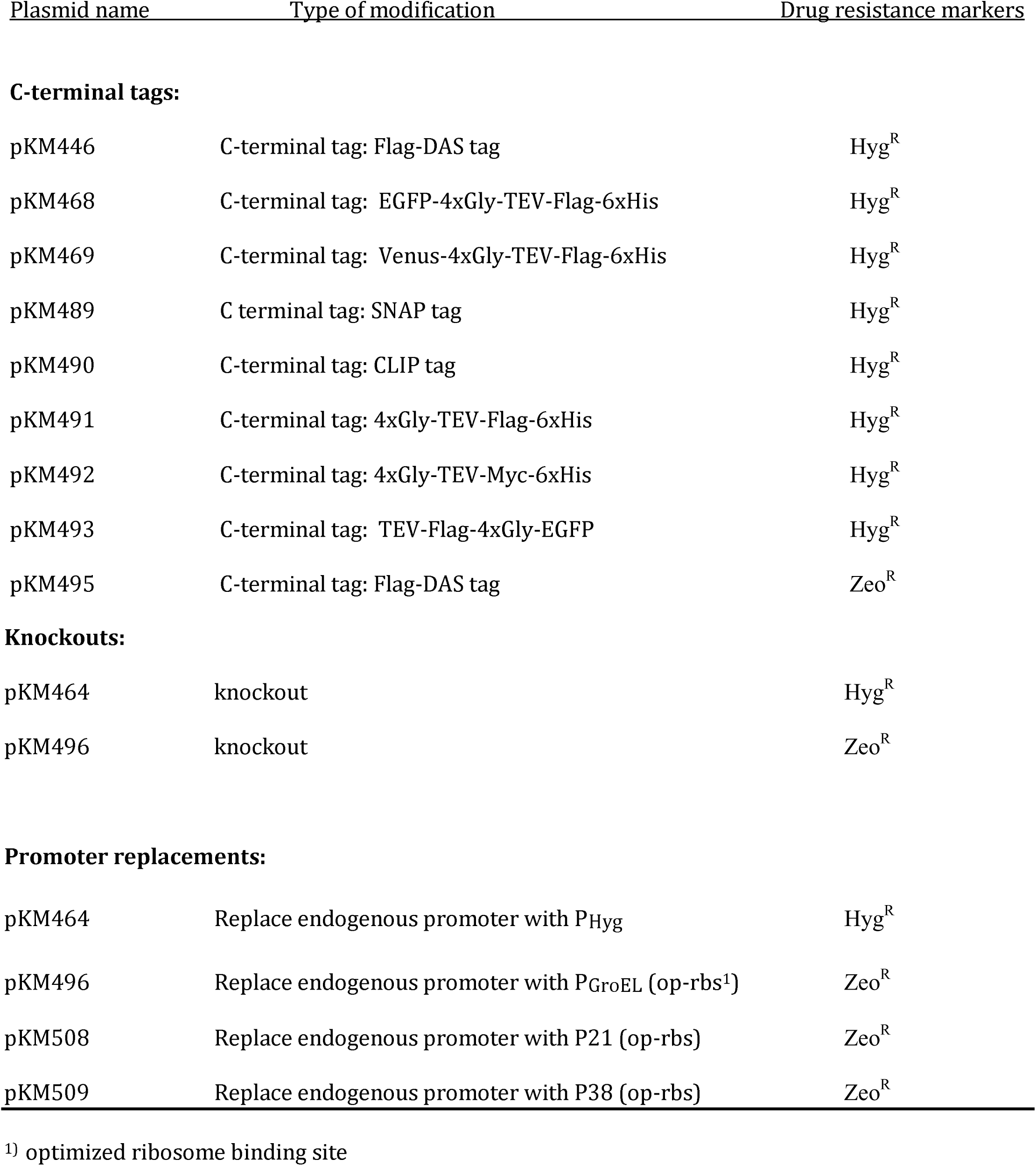
ORBIT integration plasmids.

| <u>Plasmid name</u> | <u>Type of modification</u> | <u>Drug resistance markers</u> |
| --- | --- | --- |
| **C-terminal tags:** |  |  |
| pKM446 | C-terminal tag: Flag-DAS tag | Hyg<sup>R</sup> |
| pKM468 | C-terminal tag: EGFP-4xGly-TEV-Flag-6xHis | Hyg<sup>R</sup> |
| pKM469 | C-terminal tag: Venus-4xGly-TEV-Flag-6xHis | Hyg<sup>R</sup> |
| pKM489 | C terminal tag: SNAP tag | Hyg<sup>R</sup> |
| pKM490 | C-terminal tag: CLIP tag | Hyg<sup>R</sup> |
| pKM491 | C-terminal tag: 4xGly-TEV-Flag-6xHis | Hyg<sup>R</sup> |
| pKM492 | C-terminal tag: 4xGly-TEV-Myc-6xHis | Hyg<sup>R</sup> |
| pKM493 | C-terminal tag: TEV-Flag-4xGly-EGFP | Hyg<sup>R</sup> |
| pKM495 | C-terminal tag: Flag-DAS tag | Zeo<sup>R</sup> |
| **Knockouts:** |  |  |
| pKM464 | knockout | Hyg<sup>R</sup> |
| pKM496 | knockout | Zeo<sup>R</sup> |
| **Promoter replacements:** |  |  |
| pKM464 | Replace endogenous promoter with P<sub>Hyg</sub> | Hyg<sup>R</sup> |
| pKM496 | Replace endogenous promoter with P<sub>GroEL</sub> (op-rbs<sup>1</sup>) | Zeo<sup>R</sup> |
| pKM508 | Replace endogenous promoter with P21 (op-rbs) | Zeo<sup>R</sup> |
| pKM509 | Replace endogenous promoter with P38 (op-rbs) | Zeo<sup>R</sup> |

<sup>1)</sup> optimized ribosome binding site

As with any chromosomal engineering method, one must consider the effect of the modification on expression of downstream functions, especially if the target gene is within an operon. The ORBIT knockout plasmid pKM464 was designed so that once integrated, the *hyg* gene is positioned at the junction of the insertion site allowing downstream genes to be transcribed by the *hyg* promoter. In our experience, this is generally sufficient for the generation of most mutations (see Table 1 for knockouts generated in *M. smegmatis*). For additional options, variants of pKM446 (for Flag-DAS+4 tags) or pKM464 (for knockouts) are designed that contain a second promoter (P_GroEL_) to drive higher transcription downstream of the insertion site, if needed (Table 3). Similar plasmids can be generated with promoters of varying strengths, as needed.

### GFP fusions

To demonstrate the functionality of these additional tags, ORBIT was used to construct eGFP fusions with a number of *M. smegmatis* genes. We tagged MmpL3 (the essential mycolate transporter suspected to reside at the pole) and DnaN (the beta subunit of DNA polymerase III), two proteins known to be located in discrete cytoplasmic foci, and three genes with unknown distribution patterns. The eGFP payload plasmid pKM468 (Table 3) was used to tag each gene *in situ* at the endogenous chromosomal locus. MmpL3-eGFP and DnaN-eGFP were concentrated at the predicted localization site of each protein (Fig. 7). For DnaN-GFP, the punctate spots identify positions of the replication forks and are shown as occurring in non-polar regions of the cell (see Fig. 7), in accordance with previous observations ^47^ eGFP-tagged alleles of MSMEG_3596, SppA and ClpC were all found at distinct sites, which were different from the diffuse cytosolic distribution of unfused eGFP. Overall, ORBIT is a quick and efficient way to tag native genes in the chromosome with different types of tags without having to create target‐ and modification-specific dsDNA recombination substrates. Also, by expressing each gene at its native level, aberrant localization or complex formation due to overexpression can be avoided.

**Figure 7.**
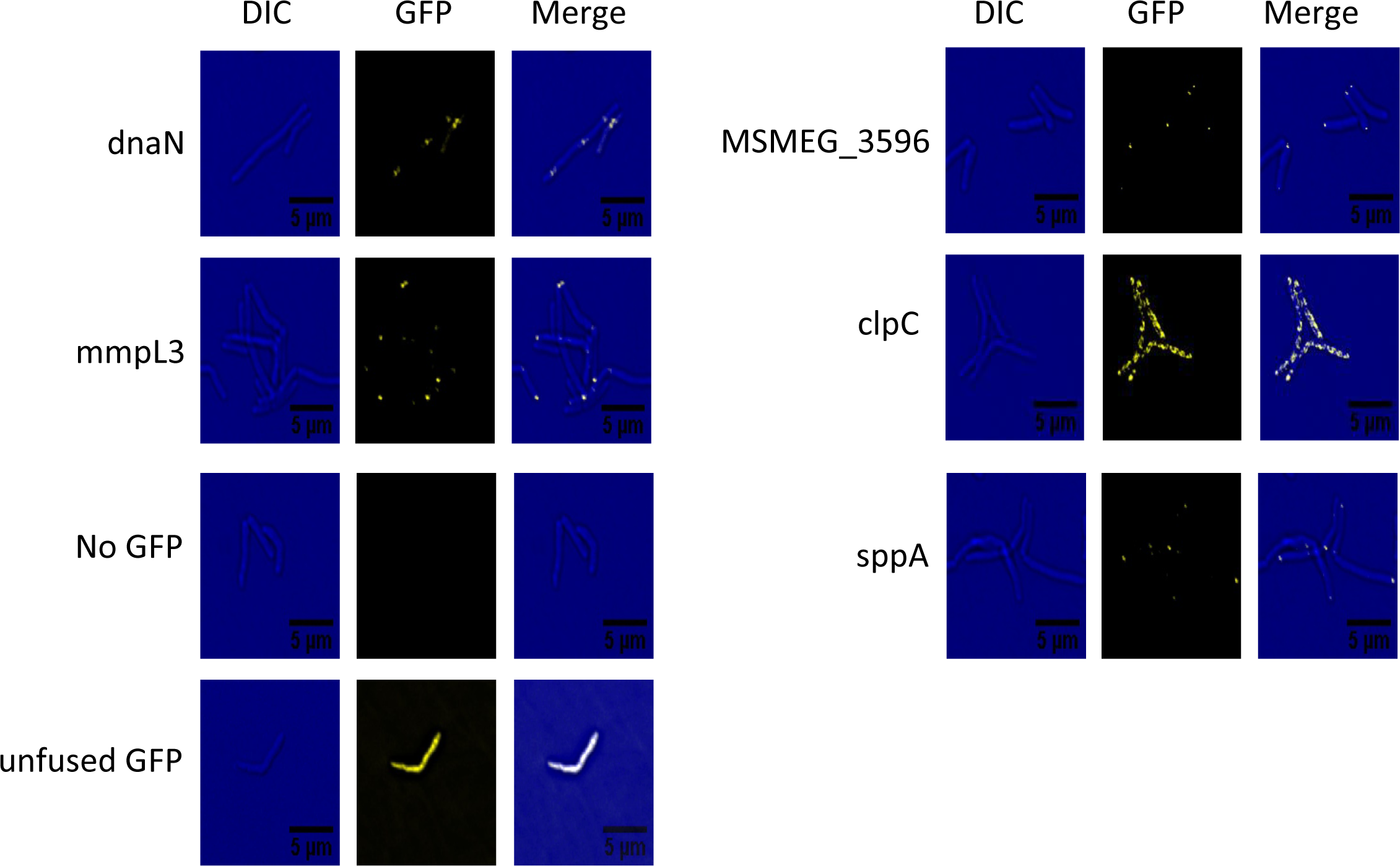
ORBIT-generated GFP fusions. *M. smegmatis* cells containing GFP-tagged targets were grown in 7H9-AD-Tween-80 to an optical density of 0.8. One microliter of the culture was spotted on an agarose pad for microscopy. Each bacterial strain was imaged using differential interference contrast (DIC) and GFP channels.

### Promoter replacements

One can also integrate *attB*-containing ORBIT plasmids to change the endogenous levels of expression of a chromosomal gene. We constructed a series of ORBIT plasmids containing promoters of different strengths that can be used to drive the expression of genes downstream of the plasmid insertion site (see Table 3). For use of these plasmids, the ORBIT oligo is designed to replace the endogenous promoter with *attP*. The target gene’s ribosome binding site, if recognized, can also be deleted, replaced, or left intact, depending on the final level of expression desired. To test this scheme, we performed ORBIT in an *M. smegmatis* strain where a chromosomal *lacZ* gene is under control of the mycobacterial PAg85 promoter containing a weak ribosome-binding site.

ORBIT was used to replace PAg85 with one of three different promoters: P_imyc_, P_GroEL_ or P38. Plasmids containing these promoters were obtained from D. Schnappinger and S. Ehrt and are listed according to increasing strengths of expression (D. Schnappinger, personal communication). The promoters were transferred to ORBIT integration plasmids and placed upstream of the Bxb1 *attB* site (see Fig. 8a). An optimized ribosome binding site (rbs: AGAAAGGAGGAAGGA) was included between the promoters and the *attB* site to increase the overall expression of β-galactosidase relative to the starting strain, where an endogenous Shine-Delgarno sequence could not be recognized. ORBIT recombinants were identified by resistance to zeocin and verified by chromosomal PCR analysis as described above. As seen in Fig. 8b, the total amount of β-galactosidase in each extract increased in accordance with the expected strengths of the three promoters used in these assays. Clearly, endogenous promoters can easily be altered using ORBIT to express different levels of a target gene.

**Figure 8.**
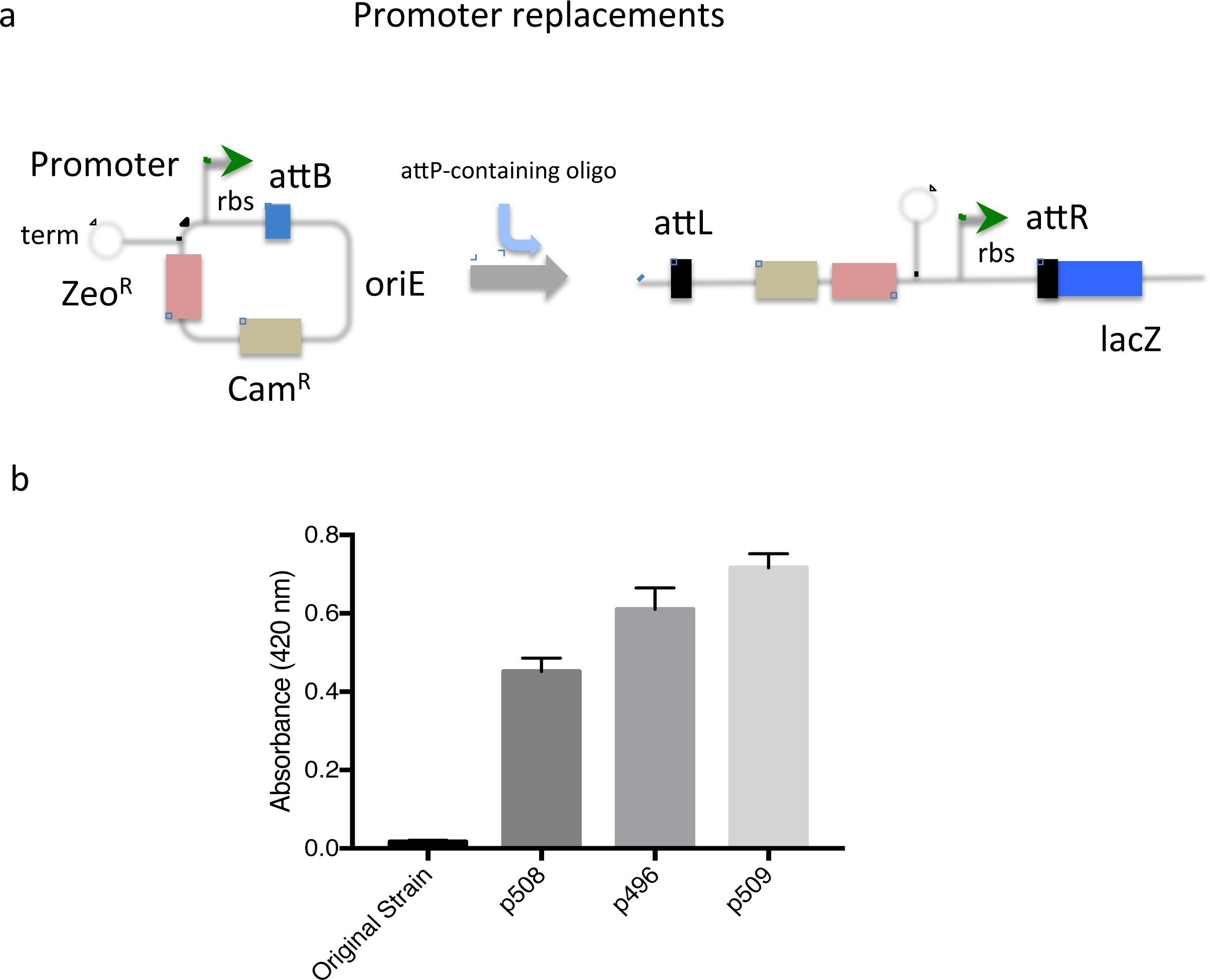
Promoter replacements. (a) Diagram of ORBIT-generated promoter replacement. In the nonreplicating plasmid, the promoter to be inserted into the chromosome is placed to the left of the *attB* site. A *TrrnB* terminator is placed upstream of this promoter to prevent read-through from the plasmid backbone. The oligo is designed to place attPjust upstream of the target gene in place of the endogenous promoter. Following integration, the promoter and inserted ribosome binding site drive expression of the chromosomal target gene (*lacZ*). (b) ORBIT was carried out with plasmids pKM496 (P_GroEL_), pKM508 (P_imyc_) and pKM509 (P38) with an oligo that deletes the endogenous promoter. Extracts of the cells were made and beta-galactosidase assays were performed in triplicate (standard error bars shown). The higher amounts of beta-galactosidase present in the engineered strains, relative to the starting strain, is likely due to the presence of the optimized ribosome binding site following each promoter.

## DISCUSSION

The λ Red recombineering system has been critical for genome engineering of *E. coli* and related bacteria. While λ Red does not function well in bacteria more distantly related to *E. coli*, endogenous phage recombination systems have been identified and employed to promote recombineering in these organisms. One of the more notable examples of a phage recombination system utilized in this way is the Che9c mycobacterial RecET system, identified by Van Kessel and Hatfull and used for recombineering of the *M. smegmatis* and *M. tuberculosis* chromosomes ^14, 15^. This recombineering system has been used to construct numerous knockouts and modifications of mycobacterial targets ^18, 39,46, 48^. However, unlike λ Red recombineering in *E. coli*, this process requires the cumbersome construction of a recombineering substrate containing >500 bp of flanking homology for each specific modification. Similarly, specialized transduction also requires the laborious construction and packaging of a different recombination substrate-containing phagemid for each genome modification^55^.

ORBIT overcomes the limitations of existing methods by combining components of two different efficient recombination systems from mycobacteria: the Che9c annealase from the general homologous recombination system and the Bxb1 Integrase from the site-specific recombination system. The construction of long dsDNA recombination substrates is replaced by the synthesis of a targeting oligonucleotide, and the availability of a library of payload plasmids allows a single oligo to be used to generate a wide variety of functional modifications. The utility of ORBIT extends beyond the simple generation of mutants. Since targeting oligos are so easily generated, the system is well suited for the construction of libraries of mutants. The targeting oligo loop containing the 48 bp *attP* site can be extended to 60 bp (Fig. 1) to include a nearly infinite number of unique DNA barcode sequences, which would allow each mutant in a large ORBIT-generated pool to be monitored independently by PCR or next-generation sequencing. The variety of modifications that can be made with the existing collection of payload plasmids opens new avenues for functional screening of mutant libraries that are generated using this method. ORBIT can also be used for genome reduction strategies in MTb. The largest deletion generated in this study was the one-step 12kb deletion of the *pks12* gene, and it is likely that even larger deletions are possible. Finally, though most of the payload plasmids described in this report are designed to modify endogenous genes, one could also use this system to place exogenous elements, such as large clusters of biosynthetic genes, into any specified position in the mycobacterial chromosome.

The paradigm of ORBIT is likely to be useful in many different bacterial species. The Bxb1 Integrase requires no host functions to carry out the site-specific and directional (*attB* x *attP*) recombination reaction. It is for these reasons that Bxb1 Integrase has been selectively employed for genetic manipulations in both bacterial and mammalian cells ^49, 50^. Thus, the key to the development of ORBIT for other microbial systems would be to find a λ Beta or RecT-like annealase that promotes some level of oligo-mediated recombineering. Datta et al ^51^ have tested a number of single-stranded annealing proteins (SSAPs) from both Gram-positive and Gram-negative bacteria for oligo-mediated recombineering in *E. coli*. While the recombineering efficiency ranged over three orders of magnitude, some SSAPs from distant species worked as well as the λ Beta protein in *E. coli*, suggesting that some annealases are easily transferable between species. The annealase for use in ORBIT would have to be able to promote annealing of an oligo containing a 48 bp *attP* insertion. Note, however, that even if oligo recombineering occurs at low frequencies with such a substrate (~10^-6^), the number of *attP* sites generated could very well be adequate for the Bxb 1 Integrase to promote integration of a non-replicating plasmid, thus making integration of the oligo a selectable event.

## Methods

### Bacterial strains

*M. smegmatis* strains used in this study were derived from mc^2^155; the *M. tuberculosis* strains were all derived from H37Rv.

### Media

*M. smegmatis* was grown in Middlebrook 7H9 broth with 0.05% Tween 80, 0.2% glycerol, 0.5% BSA, 0.2% dextrose, and 0.085% NaCl; transformants were selected on LB plates (DIFCO) with appropriate drugs. *M. tuberculosis* was grown in 7H9 broth with 0.05% Tween 80, 0.2% glycerol and OADC (Beckton-Dickinson); transformants were selected on 7H10 plates with 0.5% glycerol and OADC. When needed, antibiotics were added at the following concentrations: kanamycin (20 μg/ml), streptomycin (20 μg/ml), hygromycin (50 μg/ml), zeocin (25 μg/ml).

### Plasmids

Plasmids containing the P_imyc_ promoter ^52^, the P_GroEL_ promoter and P38 promoter were obtained from D. Schnappinger and S. Ehrt, as were plasmids pGMCgS-TetOFF-18 and pGMCgS-TetON-18, where the *E. coli* SspB adapter protein is under control of the wild type and reverse TetR repressors, respectively. Plasmids constructed for this study are described in Tables 3 & 4 and will be made available from the Addgene plasmid repository site. Details of plasmid constructions are available upon request.

**Table 4.** ORBIT-testing and promotion plasmids.

| Plasmid | Functions | Drug resistance marker |
| --- | --- | --- |
| pKM433 | Phage L5 integrating vector; HygΔ60 bp internal deletion; oriE | Zeo<sup>R</sup> |
| pKM444 | P<sub>Tet</sub>-Che9c RecT-Bxb1 Int; TetR, oriE, oriM | Kan<sup>R</sup> |
| pKM461 | P<sub>Tet</sub>-Che9c RecT-Bxb1 Int; SacRB; TetR, oriE, oriM | Kan<sup>R</sup> |
| pKM511 | P<sub>Tet</sub>-Che9c RecT-Bxb1 Int; SacRB; TetR, oriE, oriM | Zeo<sup>R</sup> |

### Oligos

Oligos used for ORBIT were obtained from IDT as Ultramers at a concentration of 100 uM and delivered in 96-well plates; they were supplied desalted with no further purification. Oligos were diluted ten-fold in 10 mM Tris-HCl, pH 8.0, and final concentrations (250-350 ng/ml) were determined by Abs_260_ using a conversion factor of O.D. of 1 = 20 μg/ml oligo. ORBIT plasmids (200 ng) were mixed with 1 μg of oligo prior to electroporation

### Design of the ORBIT oligo

The sequence of oligos flanking the *attP* site used for ORBIT must be derived from the lagging strand of the replication fork. For design of an ORBIT oligo, start with a dsDNA sequence file of a target gene, starting 200 bp upstream of the initiation codon and ending 200 bp downstream of the stop codon. Insert the Bxb1 *attP* site shown in Supplementary Fig. 2 into the target sequence file for the type of modification required (*i.e*., knockout, C-terminal tag, or promoter replacement) as described in the Results section. If the transcriptional direction of the target gene is pointing toward the chromosomal origin in either replicore (green arrows in Supplementary Fig. 2), then select the top strand (5’ to 3’) of the “target sequence + attP” file as the lagging strand DNA in the oligo. If the transcriptional direction of the target gene is pointing away from the chromosomal origin in either replicore (red arrows in Supplementary Fig. 2), then select the bottom strand (5’ to 3’) of the “target sequence + attP” file as the lagging strand in the oligo. Example shown in Supplementary Fig. 2 is for *M. smegmatis*. Apply the same rules of MTb, but assume the *dif* region occurs at 2.2 Mb. For further details, see ^18^.

### ORBIT electroporations

A culture of *M. smegmatis* containing plasmid pKM444 (or pKM461) was started overnight by adding 100-150 μl of a fresh saturated stock culture to 20 ml of 7H9 media containing 20 μg/ml kanamycin in a 125 mL flask. Cells were grown on a swirling platform at 37^°^C. The next day, at an O.D. (600 nm) of 0.5, anhydrotetracycline (ATc) was added to a final concentration of 500 ng/ml. The culture was placed back on the swirling platform at 37^°^C for 2.75-3 hours until the culture O.D. was ~1.0. The culture was placed on ice (with swirling) for 10 min and then centrifuged at 4000 rpms for 10 min in a chilled centrifuge. The supernatant was removed, the cells were gently resuspended in 1 ml of 10% cold glycerol and brought up to 20 ml with 10% cold glycerol. The centrifugation and washing steps were repeated. After the second wash, the cells were collected by centrifugation and resuspended in 2 ml of 10% cold glycerol. Aliquots of electrocompetent cells (380 μl were added to sterile Eppendorf tubes containing 1 μg of an *attP*-containing oligo and 200 ng of an *attB*-containing plasmid (except where noted otherwise in figure legends). The cells and DNA were mixed by pipetting and transferred to ice-cooled electroporation cuvettes (0.2 cm). The cells were shocked with an electroporator at settings of 2.5 kV, 1000 OHMs and 25 μF. Following electroporation, the cells were resuspended in 2 ml 7H9 media and rolled at 37^°^C overnight. The following day, two 0.5 ml portions of the culture were spread on LB plates containing 50 μg/ml hygromycin or 25 μg/ml zeocin. Recombinant colonies were picked into 2 ml 7H9 media containing 50 μg/ml hygromycin or 25 μg/ml zeocin and grown overnight at 37^°^C. Control electroporations with no DNA were also performed.

Electroporations with *M. tuberculosis* (MTb) was done in a similar manner, with the following modifications. Cells containing pKM444 (or pKM461) were grown in 30 ml 7H9 media containing OADC, 0.2% glycerol, 0.05% Tween-80, and 20 μg/ml kanamycin. At an O.D. of ~ 0.8, ATc was added to the culture to a final concentration of 500 ng/ml. After ~ 8 h of swirling at 37^°^C, 3 ml of 2 M glycine was added to the culture. The cells were shaken at 37^°^C overnight (16-20 total hours following induction), collected by centrifugation and processed as described above, except that all steps were performed at room temperature. Recombinant colonies were picked into 5 ml 7H9-OADC-tween containing 50 μg/ml hygromycin and grown with shaking for 4-5 days at 37^°^C.

### PCR analysis for verification of Recombinants

Recombinants were verified by PCR analysis; Taq polymerase was obtained from Denville Scientific, Inc. PCR reactions were performed in 30 μl volume and contained 125 μM dNTPs, 5% DSMO, 1 μM primers, 2 μl of an *M. smegmatis* overnight culture (or heat-inactivated MTb culture) and 0.2 μl of Taq polymerase. MTb cells (O.D. around 1.5) were heat inactivated at 85^°^C for 50 minutes prior to removal from the BSL3 lab. The PCR program consisted of an initial step of 95^°^ C for 5 minutes (to lyse the cells), thirty cycles of 30 sec at 95^°^ C, 30 sec at 58^°^ C, and 1 minute at 72^°^ C, and a final polymerization step for 5 min at 72^°^C. Correct-sized PCR fragments were generated from both junctions of the payload plasmid inserted into the chromosome (see Fig. 3c). In each case, the 5’ junction was verified by a target-specific primer and an “oriE” primer (CCTGGTATCTTTATAGTCCTGTCG); the 3’ junction was verified by a target-specific primer and a “HygC-out” primer (TGCACGGGACCAACACCTTCGTGG or GAGGAACTGGCGCAGTTCCTCTGG). In some cases, the 5’ junction PCR was verified by sequencing. Target-specific primers contained sequences at least 100 bp upstream (5’) and downstream (3’) of the chromosomal sequences flanking the *attP* site in the ORBIT oligo. For knockouts, an additional PCR was performed to verify the absence of the target gene in the recombinant.

### Fluorescence microscopy

Bacterial cells were mounted on 1% agar pads and imaged with a DeltaVision Personal DV microscope followed by deconvolution using SoftwoRx software (Applied Precision). Further processing was performed using FIJI software ^53^. Image brightness and contrast were adjusted for visibility and the files were converted to 600 dpi. Representative cells are shown from multiple images of each strain.

### Beta-galactosidase activity assay

A 5 mL of culture at 0.8 – 1.0 OD was pelleted and resuspended in 1 mL of freshly prepared Z buffer (50 mM Na_2_HPO_4_, pH 7.0, 10 mM KCl, 1 mM MgSO_4_, 50 mM β-mercaptoethanol). Cells were lysed by bead beating 4 times at 6.5 M/s for 30 seconds followed by centrifugation for 10 minutes to harvest the supernatant. Protein concentrations were measured with a Nanodrop. For the activity assay, 10 μg of protein and Z buffer (total volume of 100 μL) was added in triplicate to a 96-well microplate. The reaction was started with 20 μL of 4 mg/mL ONPG in 0.1 M, pH 7.0 sodium phosphate buffer. Once sufficient yellow color had developed, the reaction was terminated with 50 μL of 1 M sodium carbonate. Final absorbance of the sample was measured at 420 nm in a plate reader.

## ACKNOWLEDGEMENTS

We thank Angélica María Rivera and Desmond Goodwin for technical support. We thank Graham Hatfull and Julia van Kessel for the gift of pJV53, Dirk Schnappinger and Sabine Ehrt for plasmids, and with Eric Rubin, for early discussions on the concept of this technology. We also thank Michelle Bellerose, Yi Cai, and Clare Smith for sharing results of ORBIT modifications performed. This work was supported in part from the TB gift program from the Broad Institute, Cambridge Massachusetts.

---

**Supplementary Figure 1.**
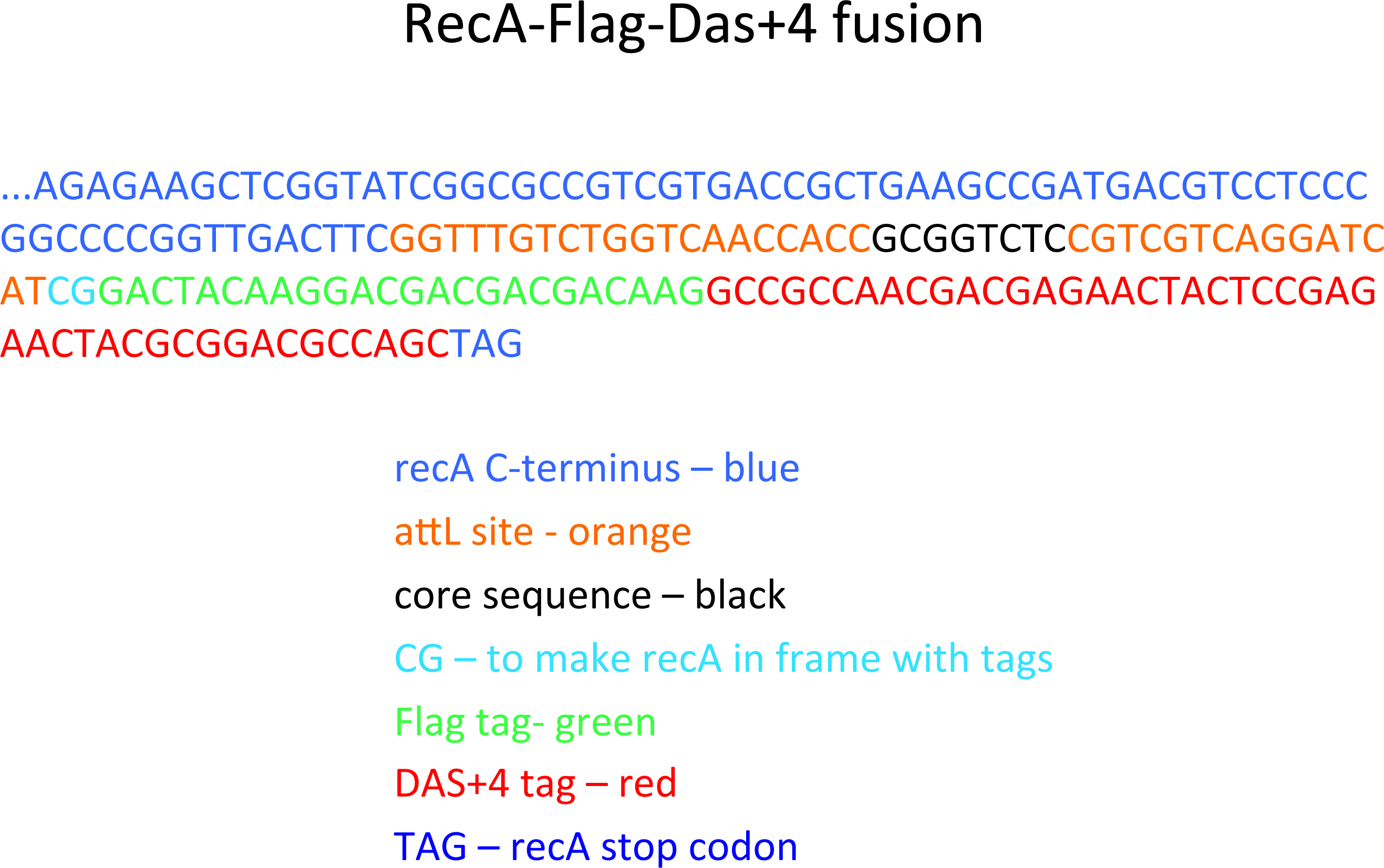
Sequence of the *rec4*-Flag-DAS+4 fusion. The sequence contains the C-terminal region of *recA* (blue), the 43 bp *attR* site (orange) created by a crossover between *attP* and *attB* (with the crossover core sequence in black), and a CG base pair (light blue) included in the ORBIT plasmid to fuse the 43 bp *attR* site with both the Flag tag (green) and the DAS+4 tag (red). Finally, the stop codon from *recA* is shown in blue.

**Supplementary Figure 2.**
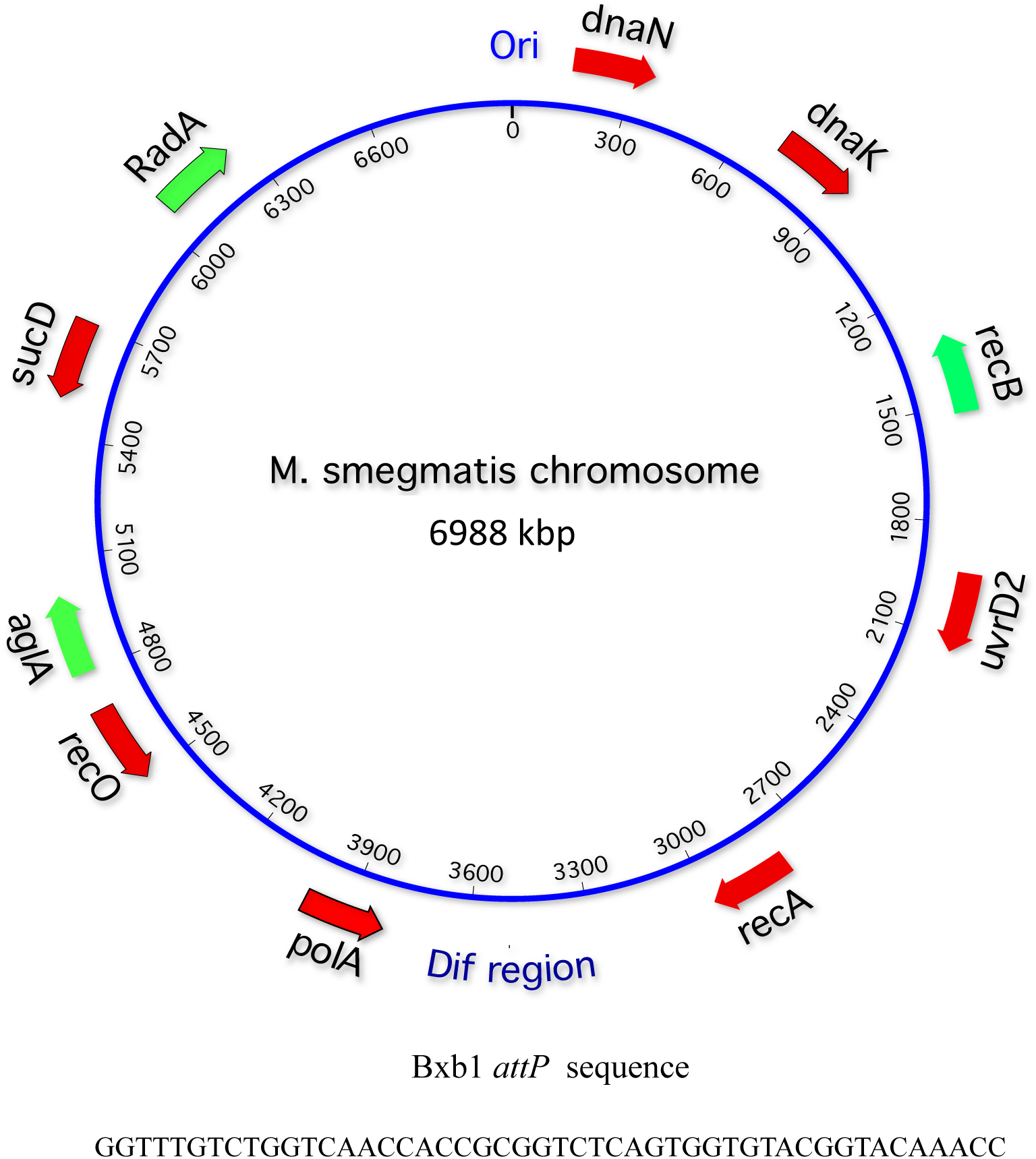
Design of the ORBIT oligo. To identify the lagging strand from a sequence file of a target gene (reading from the start codon 5’ to 3’), first insert the *attP* site into the desired position. Then, use the top strand if your target gene is transcribed toward the *ori* sequence (e.g., green arrows), or use the bottom strand if your target gene is transcribed toward the *dif* region (e.g, red arrows).

